# Microbial composition of enigmatic bird parasites: *Wolbachia* and *Spiroplasma* are the most important bacterial associates of quill mites (Acari: Syringophilidae)

**DOI:** 10.1101/377218

**Authors:** Eliza Glowska, Zuzanna K. Filutowska, Miroslawa Dabert, Michael Gerth

## Abstract

The microbiome is an integral component of many animal species, potentially affecting behaviour, physiology, and other biological properties. Despite this importance, bacterial communities remain vastly understudied in many groups of invertebrates, including mites. Quill mites (Acariformes: Syringophilidae) are a poorly known group of permanent bird ectoparasites that occupy quills of feathers and feed on bird subcutaneous tissue and fluids. Most species have strongly female biased sex ratios and it was hypothesized that this is caused by endosymbiotic bacteria. Their peculiar lifestyle further makes them potential vectors for bird diseases. Previously, *Anaplasma phagocytophilum* and a high diversity of *Wolbachia* strains were detected in quill mites via targeted PCR screens. Here, we use an unbiased 16S amplicon sequencing approach to determine other Bacteria that potentially impact quill mite biology.

We performed 16S V4 amplicon sequencing of 126 quill mite individuals from eleven species parasitizing twelve bird species (four families) of passeriform birds. In addition to *Wolbachia*, we found *Spiroplasma* as potential symbiont of quill mites. Interestingly, consistently high *Spiroplasma* titres were only found in individuals of two mite species associated with finches of the genus *Carduelis*, suggesting a history of horizontal transfers of *Spiroplasma* via the bird host. Furthermore, there was evidence for *Spiroplasma* negatively affecting *Wolbachia* titres. We found no evidence for the previously reported *Anaplasma* in quill mites, but detected the potential pathogens *Brucella* and *Bartonella* at low abundances. Other amplicon sequence variants (ASVs) could be assigned to a diverse number of bacterial taxa, including several that were previously isolated from bird skin. We observed a relatively uniform distribution of these ASVs across mite taxa and bird hosts, i.e, there was a lack of host-specificity for most detected ASVs. Further, many frequently found ASVs were assigned to taxa that show a very broad distribution with no strong prior evidence for symbiotic association with animals. We interpret these findings as evidence for a scarcity or lack of resident microbial associates (other than inherited symbionts) in quill mites, or for abundances of these taxa below our detection threshold.

## Introduction

There is abundant evidence that microbial taxa are an essential component of many animal species [1]. Bacteria-encoded traits may significantly impact host phenotypes, e.g. through providing essential nutrients [2, 3], defending against pathogens [4, 5], but also affecting ecological features of their hosts, such as mate choice [6] and life history traits [7]. Because of their potential importance in understanding the biology of many organisms, the number of microbiome studies has been soaring [8]. This popularity is owed to methodological advances (high-throughput sequencing technologies) allowing comprehensive investigation of the microbial communities [9], but also to the decreasing costs of these approaches [10]. However, the main focus of microbiome studies so far has been vertebrates [11]; while in invertebrates, the focus has been on taxa of medical, veterinary, or economical importance. For example, in mites, microbiome studies have been conducted on the pests of stored food products [12, 13], dust mites producing allergenic agents [14–16], and mites transmitting pathogens, such as sheep scab mites [17] red poultry mites [18, 19], and the honey bee parasite *Varroa* [20].

In the present study, we have focussed on quill mites (Acariformes: Syringophilidae). These obligatory bird ectoparasites live and reproduce inside the quills of feathers where they feed on subcutaneous fluids such as lymph and blood. Quill mite dispersion has been observed on the same individual (from infected to uninfected feathers), between individuals of the same species (e.g., from parents to hatchings) and occasionally by transfer between gregarious bird species [21–24]. This mode of feeding and dispersion makes quill mites potential vectors for bacterial pathogens, similar to ticks or lice [25]. However, only two bacterial taxa were recorded in quill mites so far: 1) *Anaplasma phagocytophilum* (Alphaproteobacteria, Rickettsiales) was detected in two quill mite species from three bird species [26]; 2) Multiple genetically distinct lineages of *Wolbachia* (Alphaproteobacteria, Rickettsiales) were found in five species of quill mites [27]. As these studies were targeted PCR screens, it remains unclear what other Bacteria populate quill mites. Furthermore, the importance of quill mites for bird pathogen dynamics is not known.

To address these questions, we here assess the bacterial composition of 126 quill mite individuals encompassing eleven species with a more unbiased 16S rRNA amplicon sequencing approach. We find that the symbionts *Wolbachia* and *Spiroplasma* are among the most commonly taxa associated with quill mites. Other taxa include Bacteria that were previously found in association with arthropods, and Bacteria with a very broad distribution. Strikingly, neither quill mite taxonomy nor bird host taxonomy significantly influences bacterial composition in quill mites. Furthermore, we find that despite the detection of *Bartonella* and *Brucella*, quill mites do not seem to be a major pathogen vector in birds.

## Materials and methods

### Animal collection and DNA extraction

A summary of collected quill mite species and their bird hosts can be found in Table 1. All quill mites used in this study were collected in Kopan, Poland during spring migration of birds monitored by the Bird Migration Research Station, University of Gdansk, April 2009. One secondary flight feather was completely removed from each bird specimen and dissected under a stereo microscope (Olympus ZS30). Individual mites were washed twice and preserved in 96% ethanol and total genomic DNA was extracted from single specimens using DNeasy Blood & Tissue Kit (Qiagen GmbH, Hilden, Germany) as described previously [28]. This procedure leaves the exoskeletons intact, and the specimens were subsequently mounted on microscopic slides in Faure medium, and determined using the key from Skoracki et al. (2016) [29]. All morphological observations were carried out with an Olympus BH2 microscope with differential interference contrast (DIC) optics and a camera lucida. All DNA samples and corresponding voucher specimens are deposited in the collection of the Department of Animal Morphology, Faculty of Biology, Adam Mickiewicz University in Poznań, Poland. To identify potential contaminants, in addition to sequencing a negative control alongside all samples, we further extracted DNA from reagents and materials commonly used in the laboratory this work was carried out in. One library each was created from extraction buffer (ALT), millipore water, microscope swabs, pipette swabs, and swabs of other equipment (pincettes, scalpels, benches, etc). These five libraries were processed and sequenced separately from the other samples, but by using identical procedures.

**Table 1.** Overview of quill mites sampled for this study.

| Quill mite species | Bird host species (common name) | Number of individuals |
| --- | --- | --- |
| *Aulobia cardueli* | *Carduelis flammea* (Common redpoll) | 2 |
| *Aulobia cardueli* | *Carduelis spinus* (Eurasian siskin) | 4 |
| *Syringophiloidus parapresentalis* | *Turdus iliacus* (Redwing) | 3 |
| *Syringophilopsis fringillae* | *Fringilla coelebs* (Common chaffinch) | 6 |
| *Syringophilopsis kirgizorum* | *Carduelis carduelis* (European goldfinch) | 9 |
| *Syringophilopsis kirgizorum* | *Carduelis chloris* (European greenfinch) | 2 |
| *Syringophilopsis turdi* | *Turdus iliacus* (Redwing) | 15 |
| *Syringophilopsis turdi* | *Turdus philomelos* (Song thrush) | 4 |
| *Torotrogla cardueli* | *Carduelis carduelis* (European goldfinch) | 6 |
| *Torotrogla cardueli* | *Carduelis spinus* (Eurasian siskin) | 13 |
| *Torotrogla gaudi* | *Fringilla coelebs* (Common chaffinch) | 16 |
| *Torotrogla lusciniae* | *Luscinia luscinia* (Thrush nightingale) | 7 |
| *Torotrogla lusciniae* | *Luscinia svecica* (Bluethroat) | 1 |
| *Torotrogla merulae* | *Turdus merula* (Common blackbird) | 13 |
| *Torotrogla merulae* | *Turdus philomelos* (Song thrush) | 9 |
| *Torotrogla modularis* | *Prunella modularis* (Dunnock) | 4 |
| *Torotrogla rubeculi* | *Erithacus rubecula* (European robin) | 12 |

### Library preparation and sequencing

We amplified and sequenced the V4 hypervariable region of 16S rRNA gene. For the PCRs, each 10 μl sample was prepared in two technical replicates containing 2 μl HOT FIREPol Blend Master Mix (Solis Biodyne, Tartu, Estonia), 0.25 μm of each double-indexed fusion primer (Supplementary Table S1), and about 1 ng of template DNA. The fusion PCR regime used was 12 min at 95 °C, 40 cycles of 15 sec at 95 °C, 30 sec at 58 °C, 30 sec at 72 °C, and a final 7 min at 72 °C. After PCR, all samples were pooled, size-selected on a 3% agarose gel, purified using the QIAquick Gel Extraction Kit (Qiagen), and quantified on a 2200 TapeStation (Agilent Technologies, Inc.). Clonal template amplification on Ion Sphere Particles (ISPs) was performed using the Ion Torrent One Touch System II and the Ion PGM^TM^ Hi-QTM View OT2 Kit with regard to manufacturer’s instructions. Sequencing of the templated ISPs was conducted on the Ion 318^TM^ Chip with the use of Ion PGM^TM^ Hi-Q^TM^ View Sequencing Kit and Ion PGM system (Ion Torrent, Thermo Fisher Scientific, Inc.) at Molecular Biology Techniques Laboratory, Faculty of Biology, AMU. All reads resulting from the sequencing are available under NCBI BioProject accession PRJNA482380.

### Read processing and statistical analyses

Reads were trimmed of adaptors and primer sites by using cutadapt version 1.16 [30]. The remaining reads were dereplicated, denoised, and chimeras eliminated using the DADA2 package version 1.8 [31] within the R statistical programming environment [32]. Taxonomic assignment of the ASVs (amplicon sequence variants), to species level where possible, was also performed within DADA2 using the SILVA database version 132 [33]. Next, contaminant taxa were identified from the sequenced extraction control using the ‘prevalence’ algorithm implemented in the R package decontam [34]. Further potential contaminants were identified by processing the five libraries derived from reagents and materials as described, and then excluding all ASVs that were found in any of these control libraries from subsequent analyses.

To reduce the impact of ASVs with very low abundance, we removed all ASVs that were present in only a single sample and also discarded ASVs from Bacterial Phyla that only occurred once in total. To account for potential biases between samples with uneven sequencing depth, all read counts from the remaining samples were rarefied to the read depth of the sample with the lowest read number. An overview of how our filtering steps affected ASV counts can be found in Supplementary Table S2. All subsequent statistical analyses were done on log-transformed read counts. Because the symbionts *Wolbachia* and *Spiroplasma* were dominant in some of the samples, we excluded all ASVs corresponding to these taxa prior to statistical comparisons between groups. First, we plotted the abundance of the most frequently found bacterial families using the R packages phyloseq and ggplot2 [35, 36]. Next, ordination analyses were performed with phyloseq using Bray distances and non-metric multidimensional scaling (NMDS). Differences in abundances of particular taxa between groups (quill mite species, bird host species, developmental stage, *Wolbachia* positive and negative samples) were determined with Kruskal-Wallis rank sum tests, and p-values were adjusted to these multiple comparisons to control for the false discovery rate [37]. These tests were done separately for differences in abundance of bacterial phyla, orders, families, and genera. Furthermore, we calculated Jensen-Shannon distances between the aforementioned groups and used adonis tests (analysis of variance using distance matrices) implemented in the R package vegan [38] to test if they differed significantly. The phyloseq object file containing all data used in the described analyses is available as Additional file 1.

## Results

We have investigated the microbial composition of 126 individuals belonging to eleven quill mite species that parasititze twelve bird host species of passeriform birds. Amplicon sequencing of the v4 region of the 16S rRNA gene on an IonTorrent resulted in 1,582,340 reads, with 9,426 reads per sample on average (4,616–20,231). After processing of reads (quality filtering, denoising, annotation, low-abundance filtering, rarefying, decontamination– see Materials and methods for details), 912 ASVs were retained. Among the most abundant bacterial taxa found in quill mites were *Wolbachia* and *Spiroplasma* (Fig, 1a, Supplementary Figure S1), both of which are vertically transmitted symbionts associated with a broad range of arthropods. Because these symbionts were not equally abundant across samples and might thus bias estimates of bacterial composition, they were excluded from the subsequent analyses.

**Figure 1.**
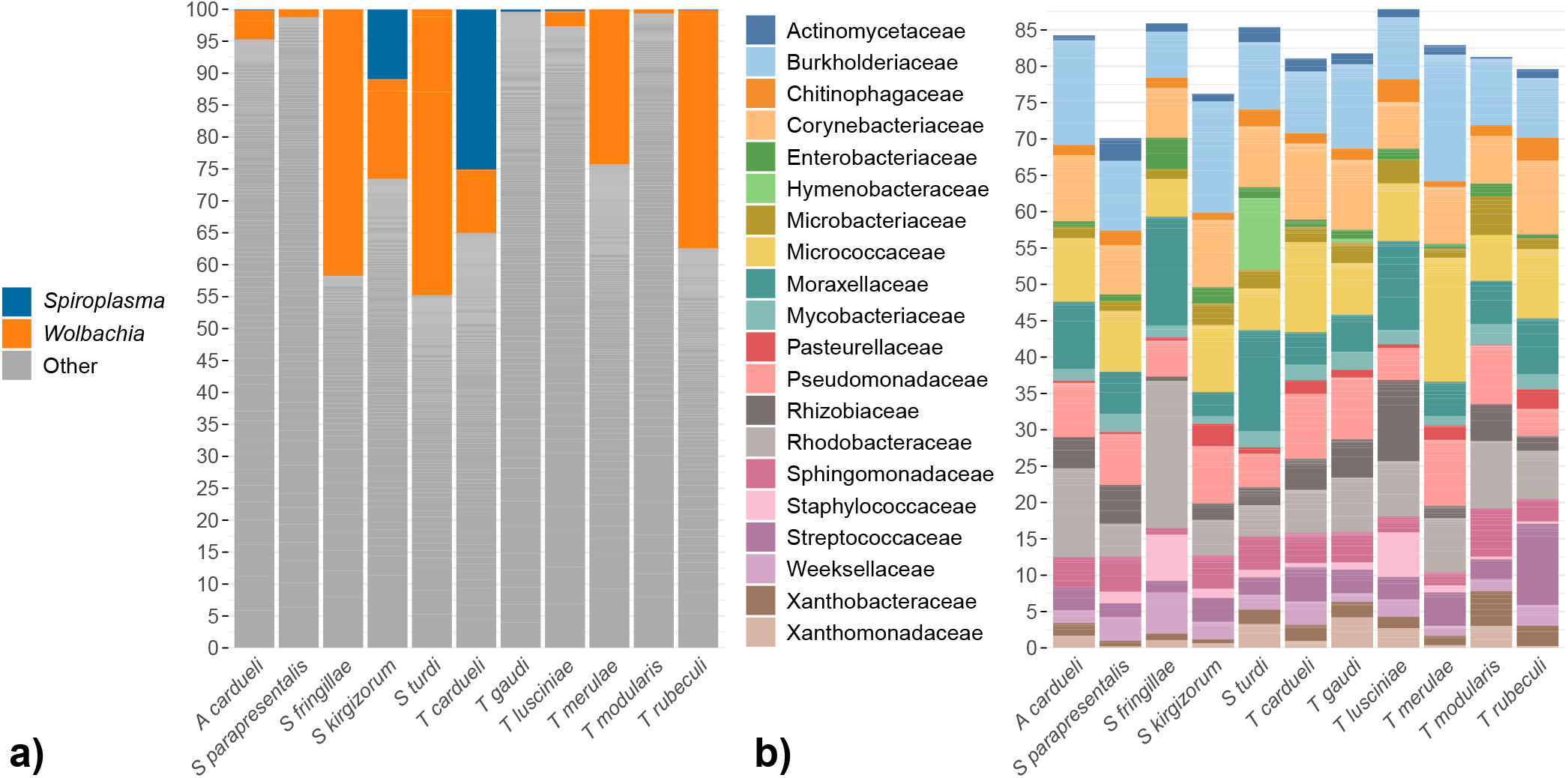
Overview of the bacterial taxa detected in quill mites. **a)** Relative abundances for the endosymbionts *Spiroplasma* and *Wolbachia*. **b)** Relative proportions of the 20 most abundantly found bacterial families in a dataset without the symbionts *Spiroplasma* and *Wolbachia*. For **a)** and **b)**, each bar represents the averaged abundances across all samples of a single species. Height of stacks represent relative abundances of each Bacterial taxon. For abundance plots of all samples, please refer to Supplementary Figure S1.

Bar plots of ASV abundance and ordination analyses with this filtered dataset revealed that the bacterial composition was relatively uniform across samples, and no clear differentiation between samples extracted from different mite species, or between *Wolbachia* positive and negative samples could be observed (Figs. 1b & 2, see also Supplementary Figure S1). However, when trying to identify differential abundance patterns of microbial composition between groups using analysis of variances, we found that bacterial composition was more similar between samples from the same quill mite species or genus and bird host species or genus than expected by chance. Furthermore, six bacterial families were found to be differentially abundant between quill mite species with a Kruskal-Wallis test (p*<*0.01, Fig. 3), one of which (Xanthobacteraceae) was also found to differ between samples of different bird host genera.

**Figure 2.**
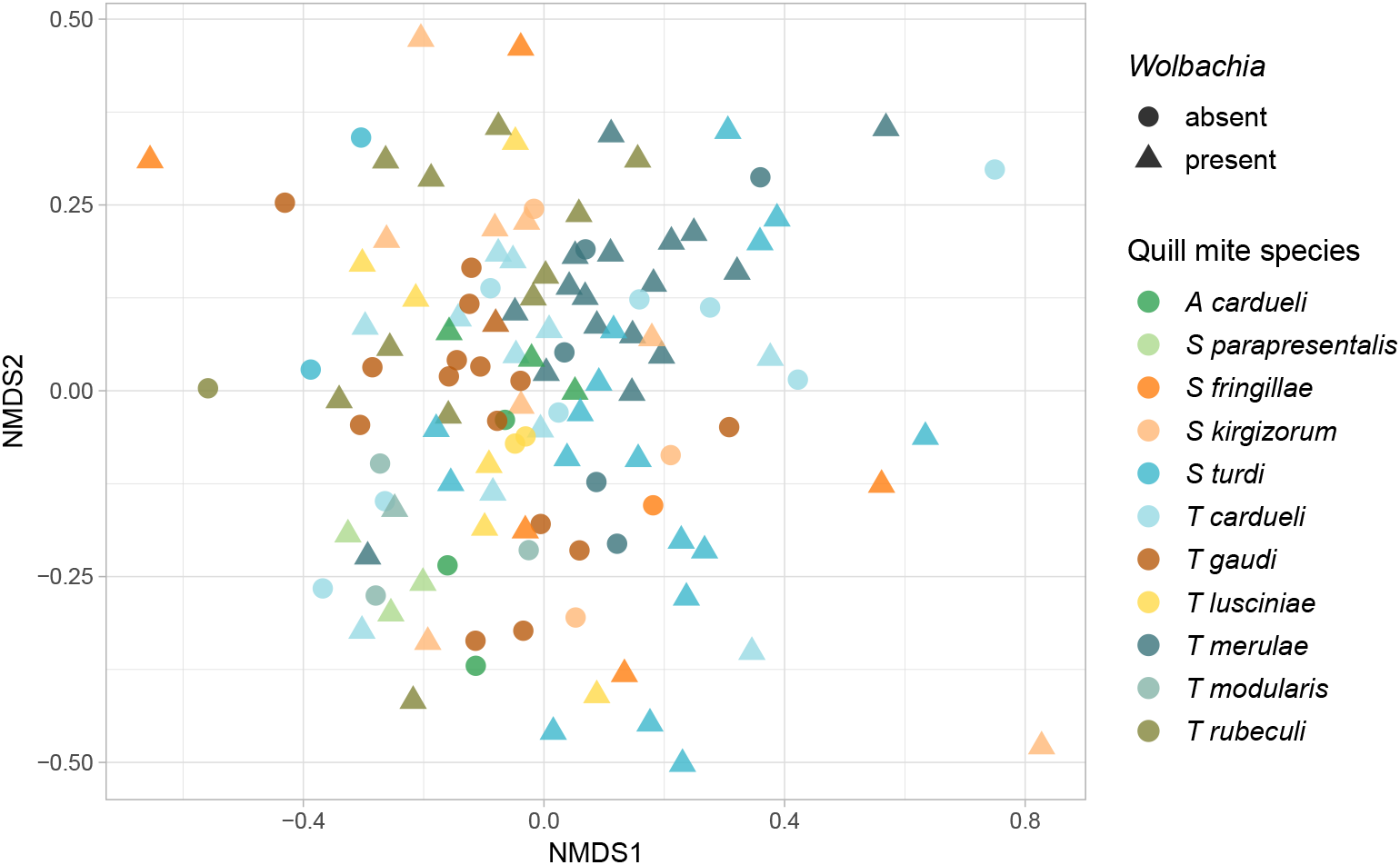
Similarity of quill mite microbiota without the endosymbionts *Spiroplasma* and *Wolbachia*. Ordination analysis is based on non-metric multidimensional scaling (NMDS) and bray distances. Log-transformed abundances were analysed. Colors of the dots represent different quill mite species from which the samples were isolated. Shape of the dots represent for *Wol-bachia* infection status.

**Figure 3.**
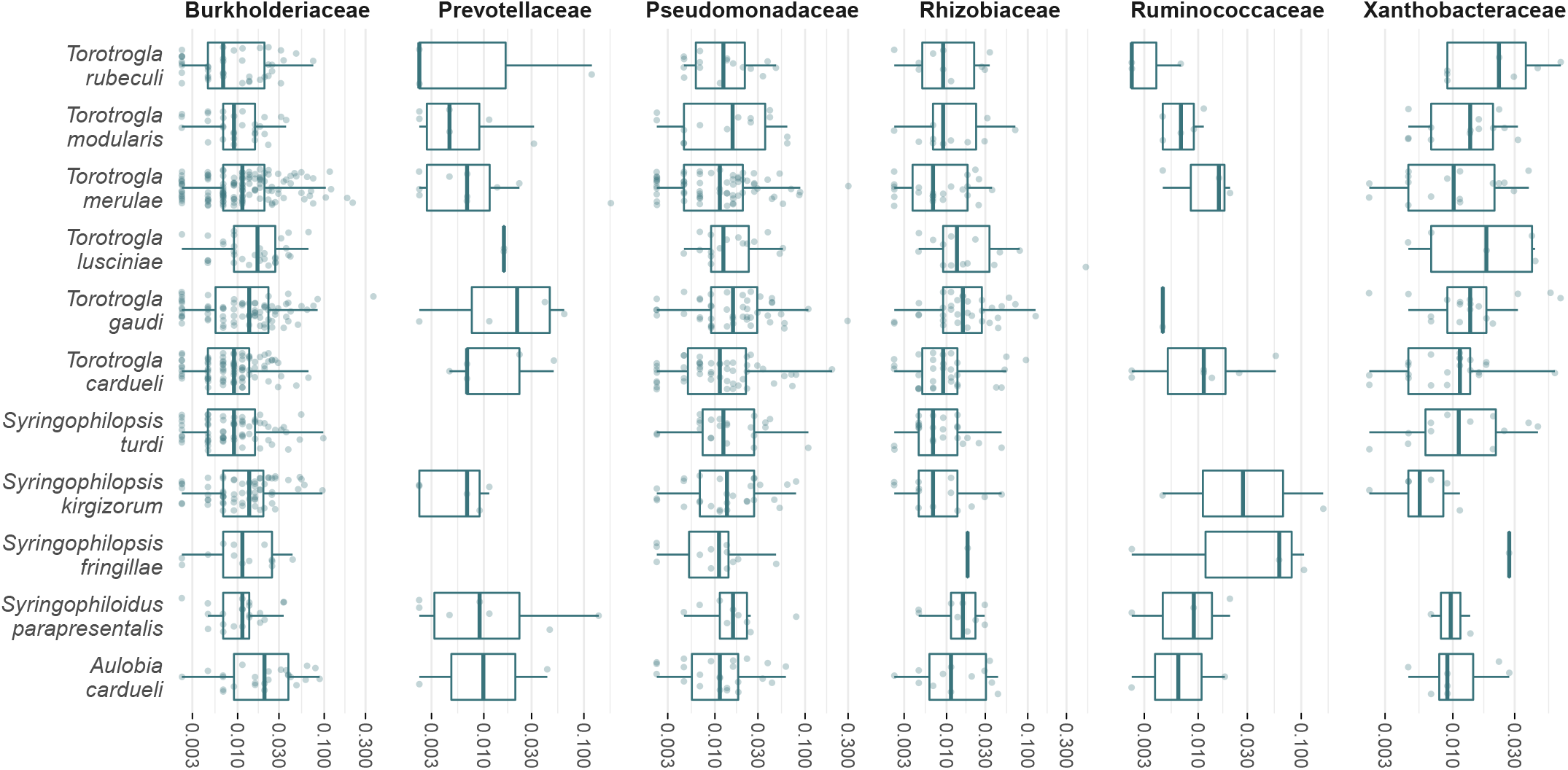
Abundance of five bacterial families that were found to be differentially abundant between quill mite species analysed. Counts for the symbionts *Wolbachia* and *Spiroplasma* were excluded.

Out of 912 detected ASVs, the ten most abundantly encountered genera were *Micrococcus*, *Corynebacterium*, *Acinetobacter*, *Streptococcus*, *Burkholderia*, *Phyllobacterium*, *Ralstonia*, *Mycobacterium*, *Paracoccus*, and *Sediminibacterium* (see Supplementary Table S2 for a full list of ASVs). None of these taxa seemed dominant in any sampled group (based on mite or bird taxonomy), and the 20 most abundant families made up similar proportions of the total ASVs across samples (Fig. 1b, Supplementary Figure S1). Other notable findings were the pathogens *Brucella* which was detected in 20 samples with an average abundance of 1.3%, and *Bartonella* which was found in two samples at 1.8% and 0.7% relative abundance, respectively.

As opposed to the general trend in the microbiome composition data, there was strong evidence for differential abundance of the symbionts *Wolbachia* and *Spiroplasma* between the bird hosts from which the mites were collected. For example, high *Spiroplasma* titres were only observed in two mite species collected from the host genus *Carduelis* (Fig. 4a, Supplementary Table S4). Further, although *Wolbachia* was present in mites sampled from all bird hosts, it was especially prevalent in mites collected from *Turdus* sp., *Erithacus* sp., and *Fringilla* sp. In contrast, it was absent or at very low titres in mites parasitizing *Luscinia* sp. (Fig. 4a). On average, the abundance of *Wolbachia* was lower in samples that also contained *Spiroplasma* (Figs. 1a, 4b). Notably, this was not an effect of *Spiroplasma* presence reducing the amount of available reads for *Wolbachia* (Fig. 4b). For mites harbouring both symbionts (eleven samples in total), we found that the abundances for *Wolbachia* and *Spiroplasma* seemed to be positively correlated (Fig. 4c).

**Figure 4.**
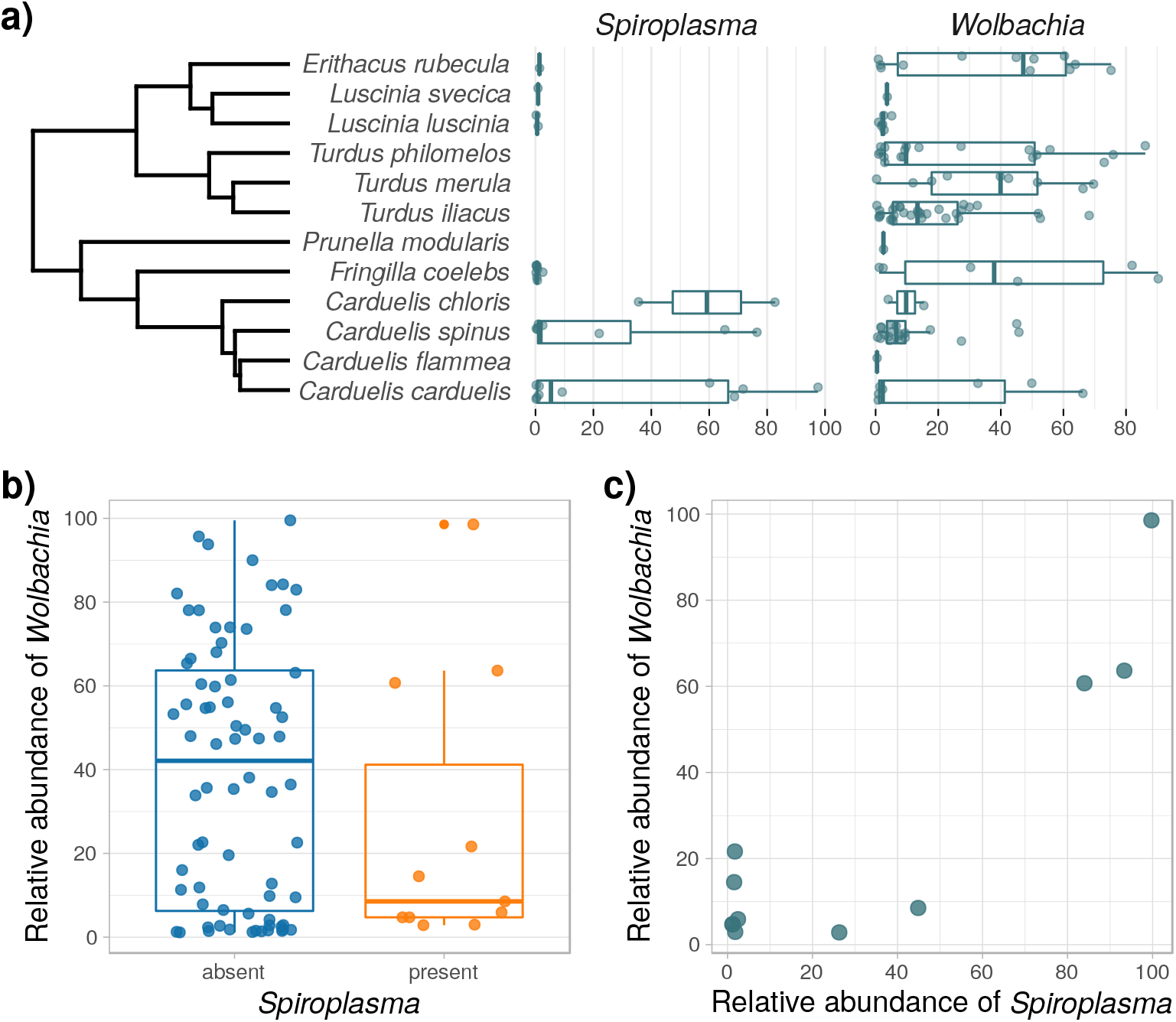
Relative abundances of the endosymbionts *Wolbachia* and *Spiroplasma* in quill mites. **a)** Abundances for all samples that are *Spiroplasma* and/or *Wolbachia* positive, sorted by bird host species from which the quill mites were isolated. Bird species phylogeny was taken from Jetz et al. ([39]; https://birdtree.org/). **b)** Relative *Wolbachia* abundances in samples with and without *Spiroplasma*. **c)** Correlation of *Wolbachia* and *Spiroplasma* abundances for samples in which both symbionts were present. For **b)** and **c)**, only samples with abundances ≥1% are shown. Also, to avoid biases of abundance estimates based on a single dominant taxon, the counts were corrected for the presence of the other endosymbiont. For example, *Wolbachia* abundance here refers to the abundance ratio of *Wolbachia* vs all other non-*Spiroplasma* ASVs. For uncorrected *Spiroplasma* and *Wolbachia* abundances for all samples, please refer to Supplementary Table S3 and Table S4.

## Discussion

### Origin of microbial DNA in quill mites

We here have sequenced microbial taxa from quill mites, an enigmatic group of bird ectoparasites. The taxa detected through 16S sequencing may be 1) resident symbionts of quill mites, 2) environmentally acquired, transient bacteria, or 3) contaminants from reagents and materials. Each of these options comes with a number of assumptions that can be tested with our data.

1. For “true”, resident symbionts, one would expect high abundances in at least some of the investigated hosts, presence in all individuals of a host species, and specialization of the symbionts, measurable as genetic differentiation between the symbionts of different host taxa. For example, all honey bees (*Apis* sp.) harbour seven core gut microbial taxa, five of which are present in other corbiculate bees, and two that are not found anywhere else [40]. The composition of these taxa is correlated with phylogenetic distances in this clade of bees, suggesting long-term association of the microbes with bees. In our dataset, *Wolbachia* and *Spiroplasma* are the most likely candidates for true symbiotic associations. Both Bacteria are known as endosymbionts from other arthropods, and are unable to permanently live outside their hosts [41, 42]. Further, we document a very high abundance of these taxa in at least some of the investigated samples (Fig. 4), which is in line with the assumptions above. In a previous study, *Wolbachia* strains of quill mites were investigated with a multi locus approach and it has been shown that quill mite associated strains are genetically very different to any other *Wolbachia* strains described so far [27]. Here, we have found 8 different ASVs annotated as *Wolbachia*, each of which is 100% identical to at least one *Wolbachia* 16S sequence previously isolated from quill mites. For *Spiroplasma*, we found a single ASV that is only 92% identical to the next closest match in the Silva database. This implies that *Spiroplasma* in quill mites might be genetically distinct from *Spiroplasma* of other arthropods, as is the case for *Wolbachia*. However, sequencing data of more loci are needed to establish the phylogenetic placement of *Spiroplasma* from quill mites.
2. For environmentally acquired, transient taxa, the expectation is that the microbial composition detected in the host reflects the microbial composition of its environment stronger than it reflects host-specific factors. For example, the gut microbiome of some caterpillars is dominated by Bacteria that derive from their food, evidenced by similar bacterial composition of leave surfaces and caterpillar faeces [43]. Quill mites live permanently within feather quills of their bird hosts, hence one might expect to find similar taxa in feathers or on bird skin as in quill mites. Unfortunately, none of the bird hosts sampled in our study was investigated previously with regard to resident skin or feather microbes. One of the most comprehensive feather microbiome studies was performed in the Dark-eyed Juncos (*Junco hyemalis*) and revealed that feathers of these birds harbour bacteria commonly occurring in the soil and phyllosphere (*Brevundimonas*, *Methylobacterium*, *Sphingomonas*), as well as potential plant pathogens (e.g. *Sphingomonas*, *Microbacterium*, *Curtobacterium*, *Rathayibacter*) [44]. All of these taxa were also found in our study, suggesting a potential environmental determinant of the bacterial composition we observed in quill mites. Furthermore, many of the core bacterial families described in bird skin microbiome studies were also found in quill mites (e.g., Pseudomonadaceae, Methylobacteriaceae, Corynebacteriaceae, Moraxellaceae, Mycobacteriaceae, Leuconostocaceae, Staphylococcaceae, Lactobacillaceae, Micrococcaceae, Streptococcaceae, Enterobacteriaceae, Sphingomonadaceae, Neisseriaceae, Xanthomonadaceae and Weeksellaceae) [45, 46]. Despite these similarities, and some statistical support for bird hosts shaping the microbiome community in our study, the lack of clustering in ordination analysis indicates that environment is not the major determining factor of quill mite microbiome composition.
3. Importantly, contaminants from reagents and kits may significantly impact microbiome compostion estimates, especially when using low biomass samples such as quill mites [47–49]. This is problematic in any microbiome study, and is very difficult to exclude with certainty. Here, we removed contaminants statistically *in silico* based on the microbial composition of the sequenced extraction control [34]. Further, we removed all ASVs present in independently sequenced controls derived from reagents and equipment commonly used in the laboratory where this study was performed (see Materials and methods). However, a number of ASVs we recovered correspond to common kit contaminants in 16S microbiome studies (e.g., *Ralstonia*, *Kocuria*), human skin Bacteria (*Corynebacterium*) or ubiquitous taxa with no strong evidence for symbiotic associations with arthropods (*Pseudomonas*, *Acinetobacter*). These taxa might constitute true associates, but we cannot exclude the possibility that they originate from contaminating sources.

In summary, we found a diverse range of Bacteria associated with quill mites. The lack of differentiation between different mite species or between species collected from different bird hosts leads us to conclude that there are no strong associations with typical gut bacteria as observed in other arthropods. However, we cannot exclude that we missed such potential associates due to the limited amount of DNA that can be extracted from the minute hosts.

### Exchange of Bacteria via bird hosts

Due to their ectoparasitic lifestyle with occasional host switching, quill mites have the potential to transmit Bacteria between their hosts. Here, we detected two pathogenic microbes that might be important in that respect: *Brucella* and *Bartonella*. *Brucella* is the agent of brucellosis, which is considered one of the most widespread zoonotic infections [50]. Several *Brucella* species are a human health threat, and people typically become infected through contact with domesticated *Brucella* infected animals, such as goats, sheep, or swine [51]. However, several bloodsucking arthropods, such as ticks and lice are regarded as possible vectors for *Brucella* [52–54]. To our knowledge, there is no data indicating that Acari other than ticks are natural *Brucella* carriers. It was hypothesized that birds and other wild animals act as natural reservoirs for *Brucella* [55], which is in line with our finding of this Bacterium in bird ectoparasites. The importance of quill mites in spreading *Brucella* between bird species remains to be assessed, but it prevalence (20/126 investigated individuals, 8 different mite species) suggests this finding is of potential importance in understanding this pathogen’s dynamics.

*Bartonella* are gram-negative Bacteria that are typically transmitted by blood sucking arthropods, and are infectious in mammalian hosts [56–58]. There are also reports on *Bartonella* incidence in birds [59, 60], and it is conceivable that the Bacteria originate from the birds, rather than from the mites. That would suggest that the host range for *Bartonella* spp. is broader than previously reported and here we expand the list of potential sources for this zoonotic infection. However, Bartonellaceae can be symbiotic in other hosts, such as honey bees and ants [61, 62]. Further, *Bartonella*-like symbionts were recently found in a number astigmatid mites [63], indicating that the *Bartonella* detected here might be quill mite symbionts, rather than pathogens. With our data, it is not possible to rule out either possibility.

Finally, we found the symbionts *Spiroplasma* and *Wolbachia* in quill mites. Both of these are common across a range of arthropod species [41, 64], are typically transmitted intraovarially, and may cause sex-ratio distorting phenotypes [65, 66]. Whereas *Spiroplasma* was so far not reported from quill mites, *Wolbachia* was previously detected and our findings confirm that this is a common symbiont of quill mites [27]. The observed presence and abundance of both taxa are not uniform across the sampled taxa (Fig. 4a). For example, *Wolbachia* is most abundant in mites parasitizing birds of the genera *Turdus*, *Erithacus*, and *Fringilla*, whereas *Spiroplasma* is most strongly associated with mites parasitizing *Carduelis*. One reason for this may be that some taxa are more susceptible than others for endosymbiosis with certain Bacteria, and this phylogenetic effect has been reported for other host taxa as well [67, 68]. Strikingly, very high *Spiroplasma* abundances were only found in two investigated mite species that are specialised parasites of three bird species of the genus *Carduelis* (Figs. 1a, 4a, Supplementary Table S4). A number of samples showed very low *Spiroplasma* titres, which may be a result of genuine low titre infections or stem from contamination via simultaneously processed libraries (e.g., through index hopping [69]). For the samples with unambiguously high *Spiroplasma* titres, the bird host phylogeny seems to be the best predictor for a *Spiroplasma* infection. One interpretation of this pattern is a history of horizontal symbiont transmission via the bird hosts. Horizontal transfers have been inferred from phylogenetic data for *Wolbachia* and *Spiroplasma* previously [70, 71], and for both symbionts, horizontal transmissions were also demonstrated experimentally [72, 73]. Although the potential mechanism of horizontal symbiont transmission via feather quills is unclear, our data suggest that the bird-parasite interactions may be important for endosymbiont transmission dynamics in quill mites.

Interestingly, we found that *Spiroplasma* presence leads to reduced *Wolbachia* titers, although this is based on a small sample size for samples that are both *Wolbachia* and *Spiroplasma* positive (N=11, Fig. 4b). Furthermore, in these eleven samples, *Spiroplasma* and *Wolbachia* titers seem to be positively correlated (Fig. 4c). It is conceivable that sharing of hosts leads to competition for finite resources the host can provide [74], and thus the growth of one symbiont might limit that of another. In *Drosophila* for example, *Spiroplasma* seem to limit the proliferation of *Wolbachia* [75] and in aphids, competition between co-occuring secondary symbionts appears to be harmful to the host [76]. Such negative fitness impacts can also expected when both symbiont titres are very high, as found here in quill mites. Although purely speculative, this may be the reason why we only observed simultaneously high *Spiroplasma* and *Wolbachia* titres in very few of the 126 investigated quill mites (Fig. 4c).

### Summary

We find a diverse, but relatively uniform set of bacterial taxa within quill mites that includes arthropod endosymbionts, pathogens, and bird associated bacteria. The importance of most of these microbes for quill mite biology is unclear, but abundances and distribution patterns suggest that *Spiroplasma* and *Wolbachia* are the most important quill mite associates.

## Acknowledgments

We thank Dr. J. K. Nowakowski, Bird Migration Research Station, University of Gdansk for the permission to collect the material. We further thank Zuzanna Filutowska for laboratory assistance. We are grateful to Piotr Łukasik for comments on an earlier version of this manuscript.

## Author contributions

Study conception and design: EG, ZKF, MG; Collecting material EG; Acquisition of data: ZKF, MD; Data analysis: MG; Interpretation of data: EG, MG; Writing the paper EG, MG. The manuscript has been read and approved by all authors.

## Supporting Information

**Figure S1**

**Overview of the bacterial taxa detected in quill mites**. **a)** Including the endosymbionts *Wolbachia* and *Spiroplasma*, and **b)** excluding these taxa. Bar plots show the 20 most abundantly found bacterial families in each dataset. Each stacked bar represents one sample, and the samples are ordered by quill mite species. Height of stacks represent relative abundances of each taxon. Note that all Anaplasmataceae ASVs are *Wolbachia*, and all Spiroplasmataceae ASVs are *Spiroplasma*.

**Table S1**

**Fusion PCR primers sequences used in this study.** Unique random barcode sequences are highlighted in bold.

**Table S2**

**Overview on the impact of filtering and decontamination on the number of retained ASVs and samples in this study.** For details on each of the steps please refer to the Materials and methods section.

**Table S3**

List of all ASVs detected in this study, ordered by abundance (relative abundances summed over all samples).

**Table S4**

Average abundance of *Spiroplasma* and *Wolbachia* across sampled bird and mite species.

**Additional file 1**

Phyloseq object including all ASVs, sample and metadata information.

